# *TFE3* fusions drive expression of CD44 and SPP1 in Translocation Renal Cell Carcinoma

**DOI:** 10.64898/2026.09.18.752493

**Authors:** Kaushal Asrani, Juhyung Woo, Thiago Vidotto, Kewen Feng, Alan Lorenzetti, Eddie Imada, Huili Li, Rajan Singh, Anand Thotakura, Sangeeta Ray, Lia Oliveira, Adrianna Amaral, Vikrant Palande, Ravi Anchoori, Jayaprakash Mandal, Roy Elias, Christopher Thoburn, Md. Masud Rana, Hrishikesh Gangule, Alexander Gao, Pornima Phatak, James Donahue, Yasser Ged, Nirmish Singla, John Copland, Laura Schmidt, W. Marston Linehan, Pedram Argani, Tamara Lotan

**Author notes:** Correspondence: Kaushal Asrani, MBBS, PhD, Department of Urology, Johns Hopkins Medicine, 600 N. Wolfe St, Baltimore, MD 21287.

## Abstract

Xp11.2 translocation RCC [Xp11.2 tRCC]) is an underdiagnosed and aggressive subtype of RCC with few specific or targeted therapies. The transmembrane glycoprotein CD44 is an emerging target in many advanced malignancies, and with its ligand OPN (*SPP1*), is a crucial driver of cancer progression, stemness, metastasis, and immune suppression. Here we show that common *TFE3*-fusions [including *SFPQ-TFE3, PRCC-TFE3, ASPSCR1-TFE3*, and *NONO-TFE3*] are associated with upregulated expression of CD44 and SPP1, as observed in multiple human and murine bulk transcriptomic studies and a murine tRCC snRNA-Seq dataset. CD44 and/or SPP1 protein expression were also upregulated in murine models of transgenic tRCC kidneys and urine specimens, patient-derived cell lines and human tRCC cases, by immunoblotting and/or IHC. Transient deletion of *CD44* was associated with profound and specific suppression of tRCC cell line growth, with decreased mTOR signaling. These data suggest that CD44 and/or SPP1 may potentially drive tumorigenesis in tRCC.

## Introduction

The MiT/TFE transcription factors (*TFEB/TFE3/MITF*) are aberrantly expressed and drive tumorigenesis in translocation renal cell carcinomas (tRCC) and a subset of potentially aggressive mesenchymal tumors called perivascular epithelioid cell tumors (PEComas)^1^. MiT/TFE proteins are critical drivers of metabolic processes; however, the specific mechanisms of tRCC tumorigenesis are not well defined^2,3^. The MiT/TFE members regulate lysosomal gene expression and drive expression of many oncogenic proteins (TGFβ, ETS-1, E-cadherin, Wnt and mTOR signaling) and cellular processes (autophagy, metabolism, cell cycle arrest), whose dysregulation is known to drive tumorigenesis. However, the molecular mechanisms underlying MiT/TFE-driven tumorigenesis are being redefined as new oncogenic transcriptional targets are discovered^4^. tRCC is a rare subtype of non-clear cell, sporadic RCC driven by chromosomal translocations involving the MiT/TFE family of transcription factors [*TFE3* (Xp11.23), *TFEB* (6p21.1), and rarely, *MITF* (3p13)]. The resulting fusions with various partner genes (e.g. *ASPSCR1, PRCC, SFPQ, NONO*), lead to constitutive nuclear localization and activation of the chimeric transcription factors^1,5^. tRCC exhibits a high degree of morphologic and clinical heterogeneity, overlapping with other RCC subtypes (ccRCC and pRCC), in part due to a diversity of *TFE3*-fusion partners with varying functions^5^, Anti-angiogenic, tyrosine kinase inhibitors (TKIs) are the mainstay of treatment for metastatic RCC (including Xp11 RCC)^6,7^. However, resistance is common and the molecular determinants of patient response have not been characterized. Moreover, tRCC is often characterized by aggressive outcomes [recurrence and metastases^8^], prompting an urgent need to identify novel biomarkers and therapeutic targets.

CD44 is a multifunctional transmembrane glycoprotein, that binds ligands like hyaluronan (HA) and Osteopontin (OPN)^9^, and mediates cell-cell and cell-matrix adhesion. CD44 regulates intracellular signaling for growth and motility, including oncogenic growth factor receptor and cytoskeletal signaling, and is a tumor-associated antigen associated with poor prognosis in many cancers. Overexpression and/or alternative splicing of CD44 have been strongly associated with tumor metastasis and progression in many tumor types^10^. CD44v6 is an important splice-variant whose expression is upregulated in many cancers including squamous cell carcinoma and adenocarcinoma^11^. Osteopontin (OPN, encoded by *SPP1*), the physiological ligand for CD44, is a multifunctional phospho-glycoprotein produced by various cell types, including immune cells and epithelial cells. OPN is overexpressed in many cancers, including ∼72% of RCCs^12^, and its expression shows strong correlation with tumor stage and aggressive phenotypes in multiple tumor types^12–14^. Importantly, SPP1 is also an immunomodulator, impacting immune cells at many levels, and the SPP1-CD44 axis can function as an immune checkpoint in multiple tumor types, negatively regulating T-cell activation in the TME to promote cancer progression^15–17^. In Tuberous Sclerosis Complex (TSC), both, CD44 (standard full-length form [CD44s] as well as the v6 splice variant [CD44v6]) and SPP1 are upregulated in pulmonary LAM driven by mTORC1 activation^18^, and CD44 is also a hallmark biomarker of perivascular epithelioid tumors (PEComas)^19,20^. In MiT/TFE tumors, mTORC1 activation has been well documented in multiple studies^4,7,8,21^, suggesting that these tumors may also demonstrated upregulation of CD44 and/or SPP1. However, the biological role of CD44 or SPP1 or their utility as biomarkers or therapeutic targets in tRCC has not been characterized.

## Results

We recently generated a transgenic mouse expressing the human *SFPQ-TFE3* fusion, downstream of a *LoxP-Stop-LoxP (LSL)* cassette (*SFPQ-TFE3^LSL^*mice)^21^. To conditionally express *SFPQ-TFE3* in tubular epithelial cells during renal development, we crossed *SFPQ-TFE3^LSL^* mice with Ksp-Cadherin (Cadherin-16 [Cdh16]) -Cre mice (***STK***). These mice developed grossly enlarged and cystic kidneys by day 15, with renal failure (elevated BUN and serum creatinine) and early neonatal death^21^. To examine effects of SFPQ-TFE3 expression following completion of kidney development, we leveraged a conditional, tamoxifen-inducible PAX8 Cre-ER^T^^2^ model^22^. At 3-4 months following tamoxifen, S*FPQ-TFE3* Pax8 ERT-Cre (***STP***) mice showed visibly enlarged kidneys with diffusely infiltrative, bilateral solid tumors, strong and diffuse, nuclear TFE3 induction, and induction of canonical MiT/TFE transcriptional targets (GPNMB, PMEL, CTSK)^21^. Expression of CD44 and its ligand SPP1, was significantly increased in bulk RNA-Seq data from the *STK* (**Fig. 1A)** and *STP* (**Fig. 1B),** transgenic mice, compared to controls. We then examined CD44 and SPP1 gene expression from RNA Sequencing data sets in murine transgenic models of tRCC and human tRCC: *CD44* and *SPP1* gene expression were increased in murine renal tumors: **a)** in the previously characterized *PRCC-TFE3; KSP-Cre* (***PTK***) mice at 7 months (**Fig. 1C)**^23^, and **b)** in the recently characterized, *ASPSCR1-TFE3; Sglt2-Cre* mice (**Fig. 1D)**^7^. *CD44* and/or *SPP1* expression was also increased in **a)** human tRCC compared to normal kidney from patients in the UTSW pan-RCC cohort [Supplemental Table **S11** from^7^], where *CD44* was one of 747 upregulated genes conserved between human and mouse tRCC [Supplemental Table **S12** from^7^, **Fig. 1E**], **b)** human fusion-*TFE3* specimens compared to normal from patients in GSE150474 (*Wang et al*)^24^ (**Fig. 1F)**, **c)** human tRCC specimens compared to normal kidney from patients in GSE167573 (*Sun et al*)^4^ (**Fig. 1G)** and **d)** *Qu et al* [Supplemental Table **S11** from ^8^]. *SPP1* also showed strong correlation with the canonical tRCC marker *GPNMB*^25^ in human tRCC cases in GSE167573(**Fig. 1H).** We also compared *SPP1* and *CD44* expression in the TCGA, comparing *TFE3* fusion-RCC and *TFEB*-amplified RCC to normal kidneys, KIRP, KIRC and KIRP cases. *SPP1* expression was significantly increased compared to normal kidney, and comparable to expression in the other cohorts (**Fig. 1I).** Finally, *SPP1* gene expression was increased in HEK293 cells with both, constitutive^3^ (**Fig. 1J)** and doxycycline-inducible^21^ (**Fig. 1K)** expression of TFE3 fusions, and in UOK cells expressing TFE3 fusions compared to normal kidney and ccRCC cell lines (**Fig. 1L).**

**Figure 1:**
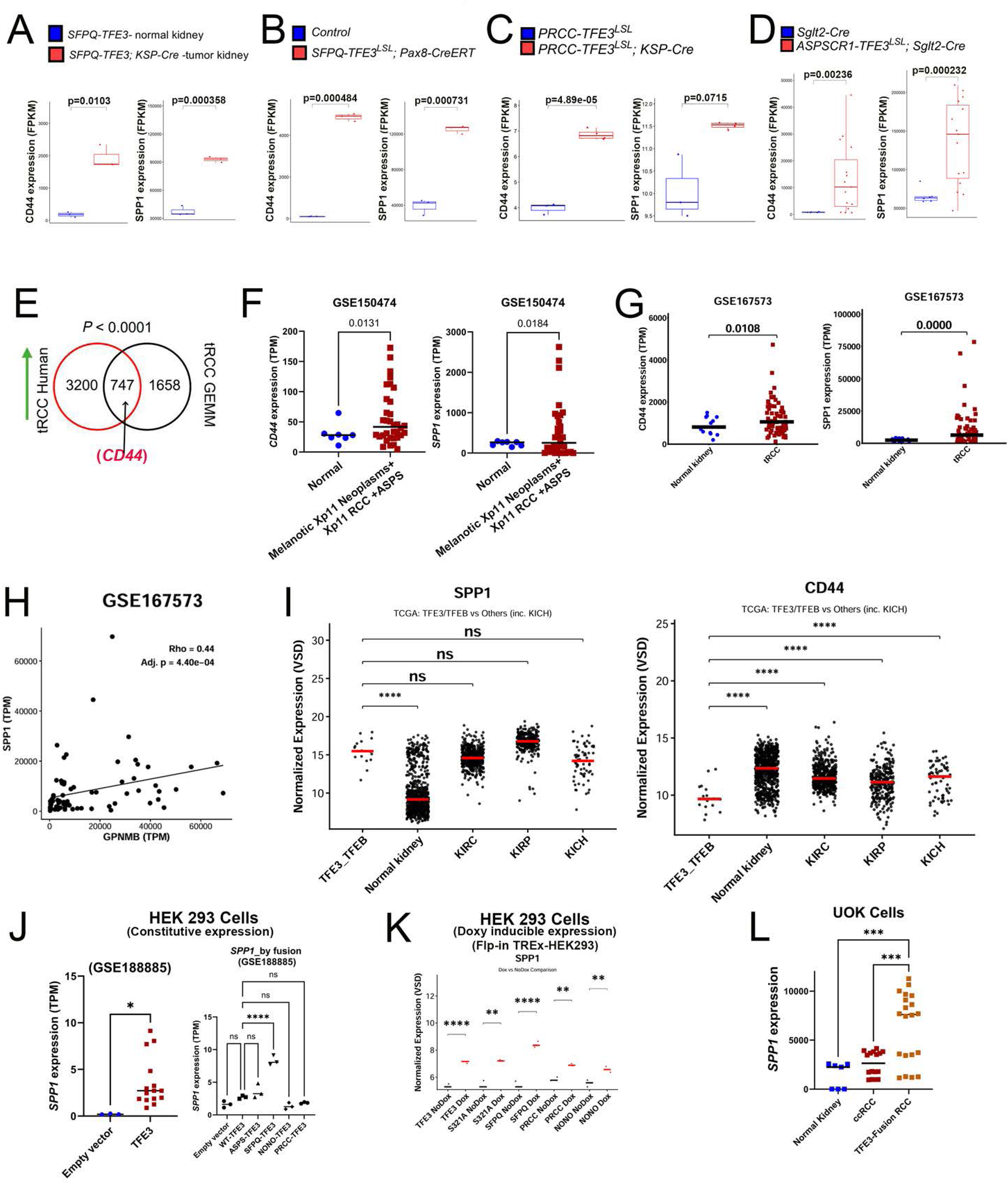
CD44 and SPP1 gene expression is upregulated in murine and human tRCC RNA sequencing datasets. **(A)** Comparison of CD44 and SPP1 expression (FPKM) in RNA seq data from: **(A)** control and SFPQ-TFE3^LSL^; Ksp-Cre transgenic mice (at day 15) from Asrani and Amaral et al. (21), **(B)** tamoxifen-injected, control and SFPQ-TFE3^LSL^; Pax8-CreERT transgenic mice at 4 months of tamoxifen injection from Asrani and Amaral et al. (21), **(C)** PRCC-TFE3^LSL^; KSP-Cre transgenic mice at 7 months, compared to controls from Baba et al. (25). **(D)** murine renal tumors in ASPSCR1-TFE3^LSL^; Sglt2-Cre mice, compared to controls from Prakasam et al. (7). **(E)** Venn diagram of significantly upregulated genes in human and murine tRCC; *CD44* is one of 747 upregulated genes conserved between human and mouse tRCC. Figure adapted from *Prakasam et al.* (7). **(F)** *CD44* (left panel) and *SPP1* expression (right panel) in GSE150474 *TFE3*-fusion cases compared to normal. **(G)** *CD44* (left panel) and *SPP1* expression (right panel) in GSE167573 tRCC cases compared to normal. **(H)** Spearman correlation showing association of *SPP1* and *GPNMB* expression in GSE167573 tRCC cases. **(I)** *SPP1* and *CD44* expression in *TFE3* fusion-RCC and *TFEB*-amplified RCC to normal kidneys, KIRP, KIRC and KIRP cases, in the TCGA. **(J)** *SPP1* expression in GSE188885, in HEK293 cells with constitutive expression of TFE3 fusions stratified by empty vs all TFE3 expressing cells (left panel) or by fusion subtype (right panel). **(K)** Comparison of *SPP1* expression (FPKM) in RNA seq data from HEK293 cells with doxycycline-inducible expression of *WT-TFE3*, *S321A-TFE3*, *SFPQ-TFE3, PRCC-TFE3* and *NONO-TFE3,* using the Flp-In-T-Rex^TM^ system from *Asrani and Amaral et al.* (21). **(L)** Comparison of *SPP1* expression (FPKM) in RNA seq data from normal kidney cells (HEK293, HK2), ccRCC cells (UOK111, UOK140, UOK150) or TFE3-fusion RCC cells [UOK120, UOK124 and UOK146 (*PRCC-TFE3*), UOK109 (*NONO-TFE3*) and UOK145 (*SFPQ-TFE3*)].

By immunoblotting, CD44 and OPN expression were increased in tamoxifen-treated, *STP* transgenic mice following 4 months of tamoxifen (**Fig. 2A),** compared to controls, although there was some variation in expression in *STP* mice, based on tumor burden. Importantly, CD44 expression was elevated and membrane-localized in tamoxifen-treated *STP* mice by IHC, with negligible expression in normal kidneys (**Fig. 2B).** Expression of OPN was also considerably elevated in *STK* (**Fig. 2C),** and *STP* (**Fig. 2D),** transgenic mice by IHC, with negligible expression in normal kidneys. We then characterized expression of CD44/OPN in the *PRCC-TFE3; KSP-Cre* (*PTK*) mice at 7 months (mice from NCI; kind gift of Dr. W. Marston Linehan)^23^. By immunoblotting, CD44 and OPN expressions were increased in *PTK* transgenic mice compared to controls (**Fig. 2E).** CD44 expression was also elevated and significantly membrane-localized in *PTK* mice by IHC, with negligible expression in normal kidneys (**Fig. 2F).**

**Figure 2:**
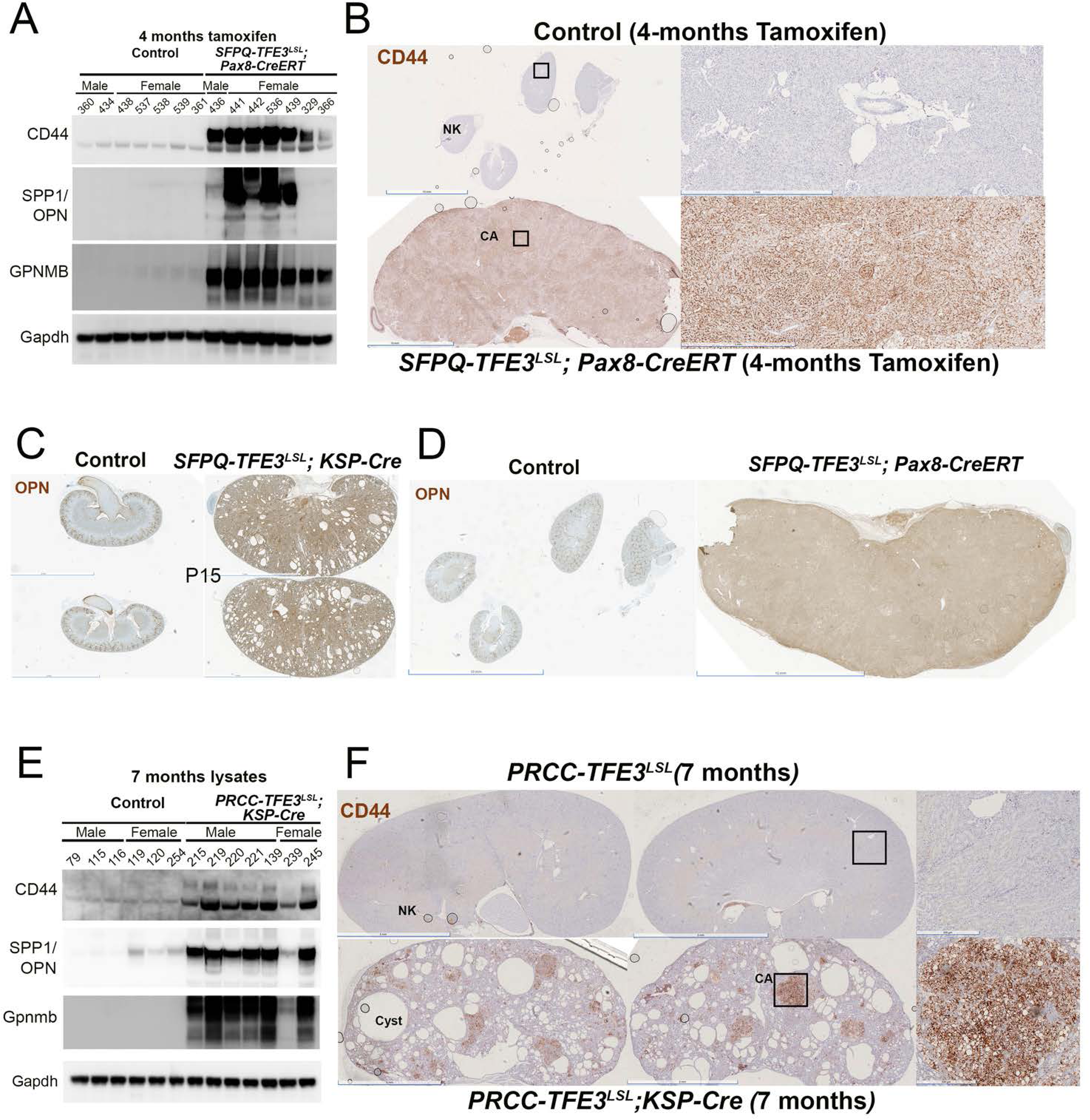
***CD44 and SPP1 protein expression is upregulated in murine models of tRCC.*** Immunoblotting of kidney lysates from **(A)** Tamoxifen-injected, control and *SFPQ-TFE3^LSL^; Pax8-CreERT* mice for CD44, OPN and GPNMB. **(B)** IHC for CD44 (37259; Cell Signaling) on tamoxifen-injected, *SFPQ-TFE3^LSL^; Pax8-CreERT* mice (bottom panels) and age-matched, littermate controls (top panels). Left panels= low magnification (scale bar=10 mm). Right panels = high magnifications from inset (scale bar=1 mm). IHC for OPN (88742; Cell Signaling) on: **(C)** age-matched controls (left panels) and *SFPQ-TFE3^LSL^; Ksp-Cre* transgenic mice (right panels) at post-natal day 15 and **(D)** tamoxifen-injected, control (left panels) and *SFPQ-TFE3^LSL^; Pax8-CreERT* mice (right panels). Scale bars=5 mm (left panels) and 10 mm (right panels). Immunoblotting of kidney lysates from **(E)** control and *PRCC-TFE3^LSL^; Ksp-Cre* transgenic mice at 7 months for CD44, OPN and GPNMB. **(F)** IHC for CD44 (37259; Cell Signaling) on 7-month, *PRCC-TFE3^LSL^; Ksp-Cre* transgenic mice (bottom panels) and age-matched, littermate controls (top panels). Left panels= low magnification (scale bar=5 mm). Right panels = high magnifications from insets (scale bar=500 µm).

To better characterize the cell-specific expression of *CD44* and *SPP1,* we performed snRNA-seq on *SFPQ-TFE3* Pax8 ERT-Cre (***STP***) mice, at both early (15 days) and late (4 months) times following induction of *SFPQ-TFE3,* as well as their respective time-matched untreated controls. Unsupervised clustering and UMAP projection of ∼ 72,904 cells identified 24 different clusters that were consolidated into 17 cell populations based on marker gene expression (**Fig. 3A-C).** The proportion of all cells were relatively consistent across the 5 samples, with the exception of the “potential cancer cell” population that was mostly absent in all the samples but strongly associated with sample B4 (SFPQ*-TFE3* Pax8 ERT-Cre: 4 months + Tamoxifen) (**Fig. 3D).** Moreover, this population also clustered separately from the remaining non-malignant cell populations based on expression of the canonical MiT/TFE transcriptional target GPNMB^23^ (**Fig. 3E).** Interestingly, *CD44* was also largely expressed in the potential tumor cluster, while expression of *SPP1* was more widespread including tumor cells but also other types (immune, proximal and distal tubules, ascending limb, etc).

**Figure 3:**
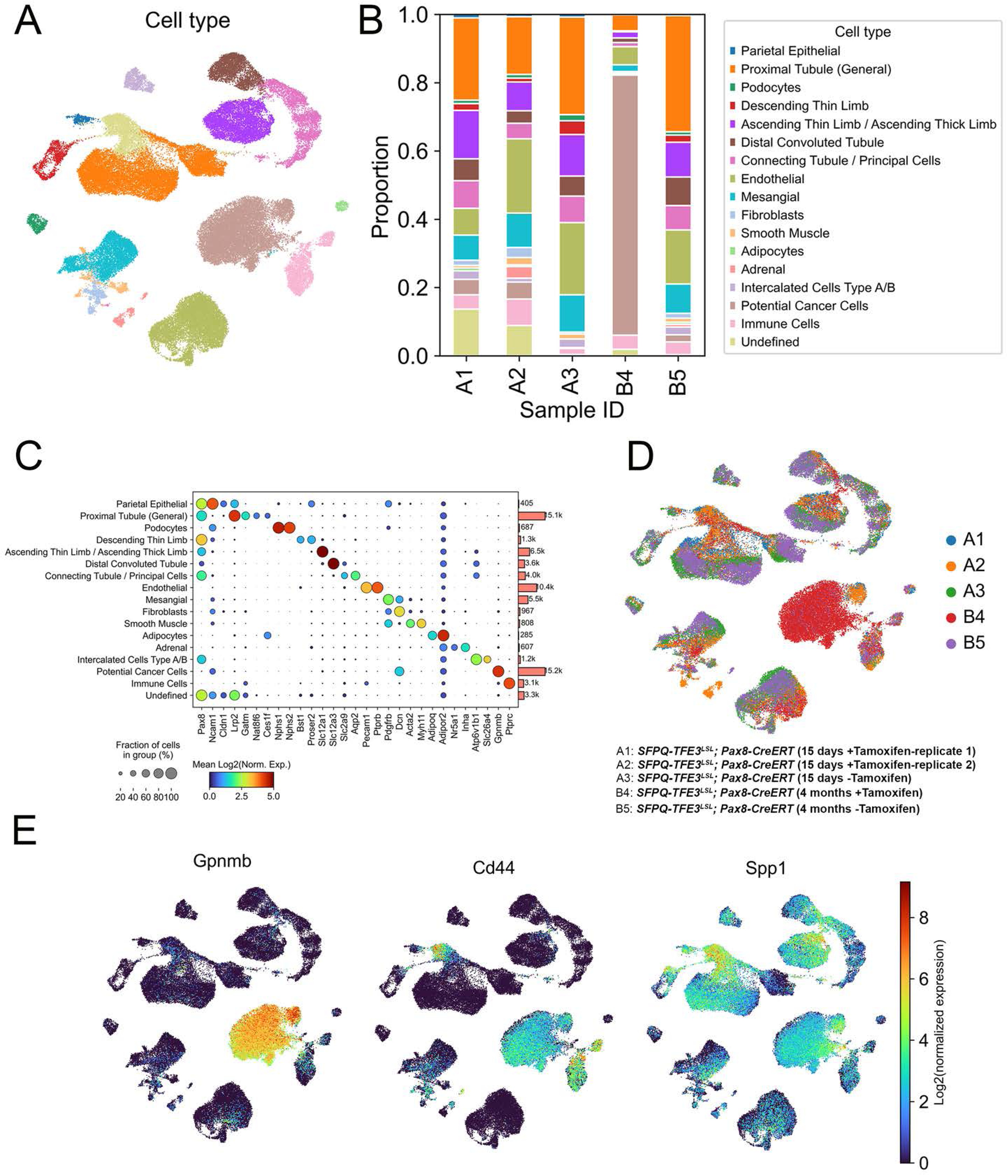
***Cell-type composition and gene expression in kidneys from SFPQ-TFE3^LSL^; Pax8-CreERT transgenic mice***. UMAP visualization of 72,904 nuclei from five transgenic mouse kidney samples, colored by **(A)** annotated cell type, and (**B)** depicting the proportion of cells of each type within the indicated samples. **(C)** Dot plot depicting the expression of kidney lineage marker genes used to classify samples into cell populations. UMAP visualization of nuclei from (A) colored by **(D)** sample id and **(E)** expression of *Gpnmb, Cd44,* and *Spp1* respectively. Each point represents one nucleus, and all panels show the same embedding computed without Harmony correction. Gene-expression colors indicate log₂ (1 + normalized expression).

We then evaluated the expression of secreted OPN, by immunoblotting of urine samples in transgenic models. In *STP* mice treated for only 2 weeks with tamoxifen and preceding development of tumors, TFE3 expression was increased in a mosaic manner (**Fig. 4A).** Strikingly, we saw a prominent increase in OPN expressions in urine samples of these *STP* mice by immunoblotting, with no expression in normal urine, suggesting that it may serve as an early and specific biomarker of renal disease (**Fig. 4B).** Urinary OPN expression was also upregulated at later points in *PTK* transgenic mice urine (**Fig. 4C).** Interestingly, OPN concentrations are also known to be elevated in plasma from LAM patients compared to normal volunteers and is a potential secreted biomarker in TSC^18^. Correspondingly, we also found that expression of CD44 and SPP1/OPN was increased in murine renal tumors with *Tsc2* loss by IHC (**Fig. 4E, F)**^26^, consistent with this prior study in LAM^18^, and potentially suggesting that upregulation of these factors may be due to increased mTORC1 activation downstream of TSC loss, or the MiT/TFE factors^26–28^.

**Figure 4:**
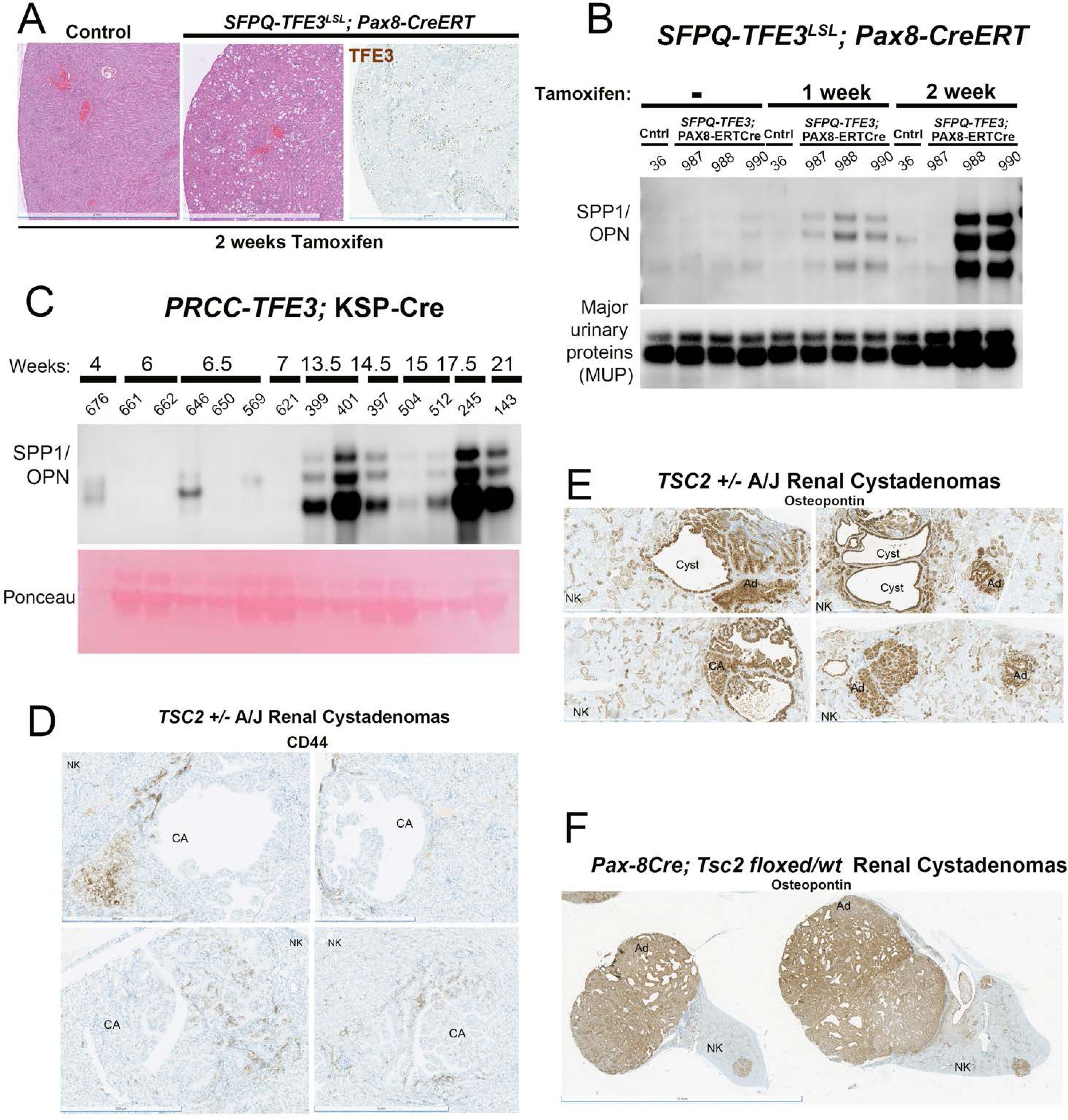
***Secreted SPP1/OPN protein expression is detected in urine samples of murine models of tRCC.* (A)** H&E and IHC for TFE3 (brown) in tamoxifen-injected, *SFPQ-TFE3^LSL^; Pax8-CreERT* transgenic mice and age-matched, littermate controls, at 2 weeks following injection of tamoxifen. Scale bar = 100 µm. **(B)** Immunoblotting shown OPN expression in urine samples of control and *STP* transgenic mice, before, and at 1or 2 weeks following treatment with tamoxifen. **(C)** Immunoblotting shown OPN expression in urine samples of *PTK* transgenic mice at different ages. **(D)** IHC for CD44 (37259; Cell Signaling) in *Tsc2 ±* murine renal cysts and cystadenomas (CA), compared to surrounding normal kidney (NK). IHC for OPN (88742; Cell Signaling) in murine renal cysts, adenomas, and cystadenomas (CA), compared to surrounding normal kidney (NK) from **(E)** *Tsc2 ± A/J* mice and **(F)** *Pax8 Cre; Tsc2fl/wt* mice.

We then examined expression of CD44 and OPN in tRCC human cell line models^29^, comparing them to ccRCC cells by immunoblotting-expression of total CD44 was increased in starved UOK124 cells bearing *PRCC-TFE3* fusions, compared to ccRCC controls (UOK111) (**Fig. 5A).** Expression of secreted SPP1/OPN was also increased in a time-dependent manner in conditioned media of UOK124 cells compared to UOK111 controls (**Fig. 5B).** In HK2 cells (human proximal renal tubular epithelial cell line) with doxycycline-inducible expression of the *PRCC-TFE3* fusion using the rtTA3 (Tet-on) construct^30^ (kind gift of Dr. Laura Schmidt, NCI), expression of the v6 splice variant [CD44v6]) was increased by immunoblotting (**Fig. 5C).** CD44 protein expression was also upregulated in primary renal epithelial cells from *SFPQ-TFE3^LSL^* mice, treated with adenoviral Cre-recombinase (**Fig. 5D).** Finally, CD44 expression was increased, and was significantly membrane-localized in FFPE specimens of xenografts derived from *PRCC-TFE3* cells [UOK124 and UOK146], compared to ccRCC [UOK111] xenografts by IHC (**Fig. 5E),** in tumors derived from an *ASPSCR1-TFE3* patient grown in NSG mice (**Fig. 5F)** and in biopsy specimens of a hepato-portal lymph node metastases [samples courtesy, of Dr John Copland, Mayo Clinic, FL] (**Fig. 5G).**

**Figure 5:**
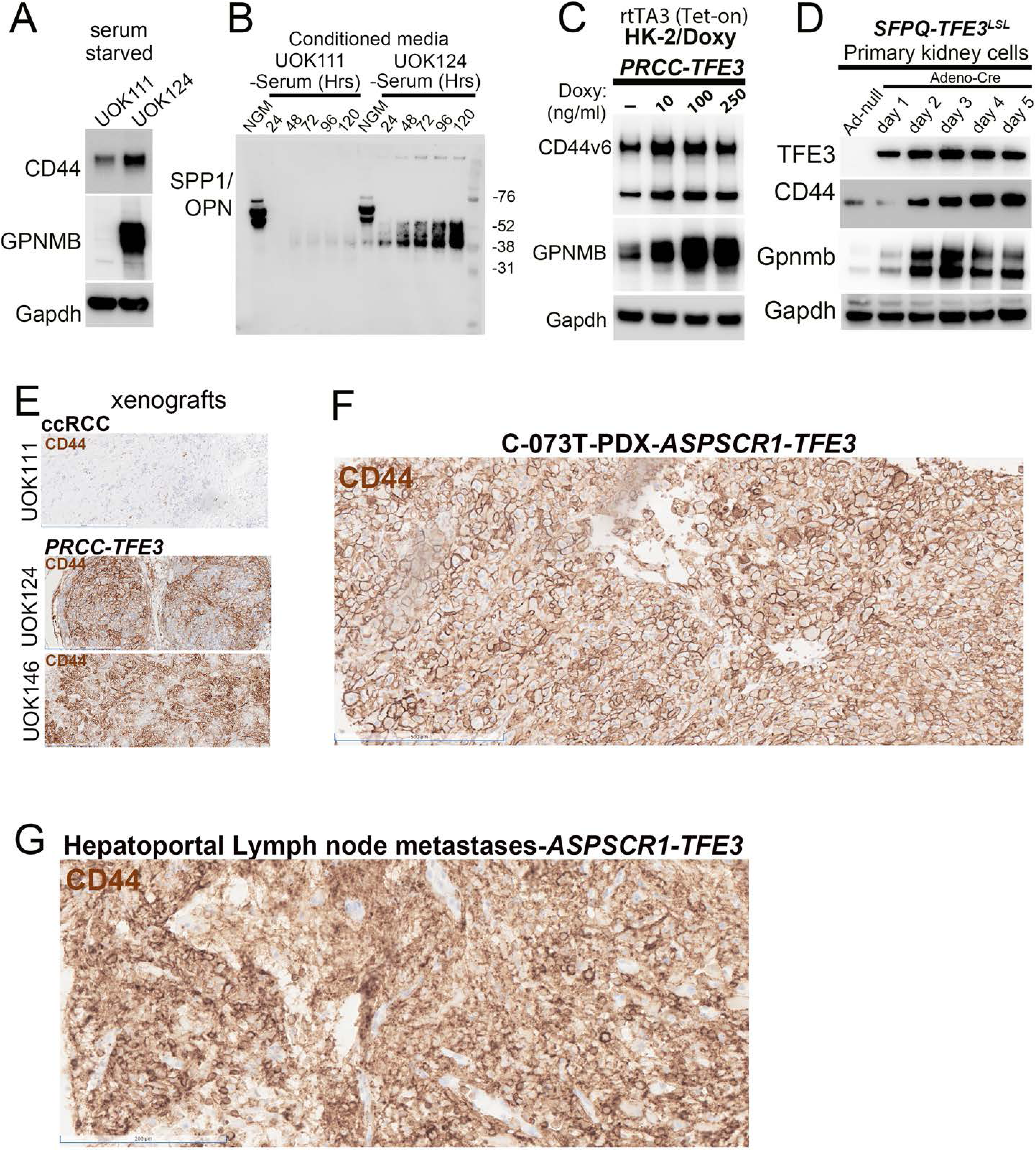
CD44 and SPP1/OPN expression is elevated in human and murine tRCC cells, human tRCC NSG xenografts, PDX mouse model and metastatic biopsies from tRCC patients. **(A)** Immunoblotting of lysates from starved, UOK124 (PRCC-TFE3) and UOK111 (ccRCC controls) cells for CD44 and GPNMB. **(B)** Immunoblotting of conditioned media from starved UOK124 and UOK111 cells, for SPP1/OPN. **(C)** Immunoblotting of lysates from HK2, doxycycline-inducible PRCC-TFE3 cells for CD44v6 and GPNMB. **(D)** Immunoblotting of lysates from primary renal tubular epithelial cells from SFPQ-TFE3^LSL^ transgenic mice treated with control or Cre-recombinase expressing adenovirus in vitro for TFE3, CD44 and GPNMB. **(E)** Representative IHC for CD44 (37259; Cell Signaling) on FFPE specimens of UOK111, UOK124 and UOK146 xenografts, performed using the Ventana Discovery ULTRA (Ventana/ Roche). Representative IHC for CD44 (37259; Cell Signaling) on FFPE tumor specimens from **(F)** an ASPSCR1-TFE3 PDX mouse model and **(G)** a hepatoportal lymph node metastatic biopsy from a patient with the ASPSCR1-TFE3 fusion-positive tumor, performed using the Ventana Discovery ULTRA (ASPSCR1-TFE3 samples were a kind gift of John A. Copland, Mayo Clinic, FL).

To characterize the effects of CD44 on cell growth, we performed transient RNAi experiments, infecting UOK124 cells with lentiviral *CD44* shRNA, which resulted in nearly complete loss of CD44 protein expression by immunoblotting (**Fig. 6A).** Strikingly, deletion of CD44 in UOK124 cells, as well as all other tRCC cell lines [UOK120, 146, 109 and 145] selectively decreased clonogenicity in crystal violet assays (**Fig. 6B, C)**, while CD44 loss in ccRCC cells [UOK111, 140, 150] had no effect (**Fig. 6D).** To further characterize these effects, we performed RNASeq on UOK124 cells with transient *CD44* deletion. By GSEA, multiple critical cell proliferation and/or cell cycle-associated genes sets [E2F targets, G2M Checkpoint, MYC Targets V1 and V2] were downregulated in the CD44-shRNA cells, consistent with the effects on growth suppression (**Fig. 6E).**

**Figure 6:**
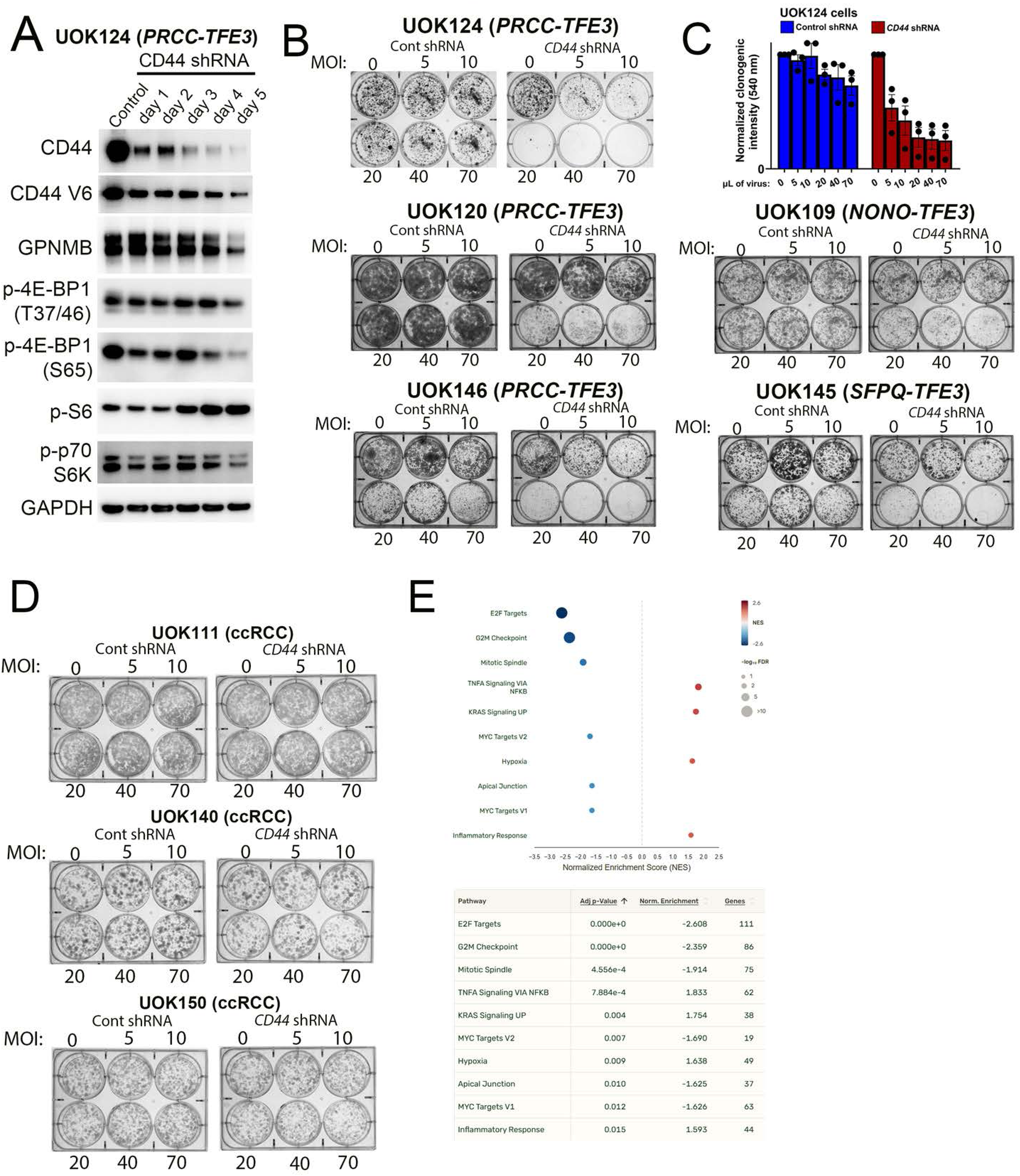
***Transient CD44 knockdown via shRNA decreases MTOR signaling and colony formation in tRCC cell lines.* (A)** Immunoblotting of lysates from UOK124 (*PRCC-TFE3*) cells transiently infected with MOI of *CD44* lentiviral shRNA for the indicated times and antibodies. **(B)** Crystal-violet stained colonies of tRCC cells [UOK124, UOK120 and UOK146 (*PRCC-TFE3*), UOK109 (*NONO-TFE3*) and UOK145 (*SFPQ-TFE3*)], following transient infection of cells with MOIs of control or *CD44* lentiviral shRNA for 10-12 days. **(C)** Normalized clonogenic intensity obtained via dissolution of colonies in 10% acetic acid and quantified using absorbance at 540nm, from UOK124 cells, infected with either control or *CD44* lentiviral shRNA for 10-12 days, from experiments in **(B)**. **(D)** Crystal-violet stained colonies of ccRCC cells (UOK111, UOK140, UOK150), following transient infection of cells with MOIs of control or *CD44* lentiviral shRNA for 10-12 days. **(E)** Functional enrichment showing significantly upregulated and downregulated pathways in UOK124 cells transiently infected with *CD44* lentiviral shRNA compared to controls, as depicted by dot plots in the upper panel. Pathways were identified with gene set enrichment analysis (GSEA) using the Hallmark gene set, following RNASeq. Adjusted p-values and Normalized Enrichment Scores (NES) of major pathways are shown in the lower panel.

## Discussion

Cumulatively, we show that CD44 and/or OPN gene and protein expression, membrane localization and/or secretion are upregulated in multiple models, and across a spectrum of *TFE3*-fusion partners, in cell line, murine transgenic and/or human renal tumors in tRCC, as well as murine models with *Tsc2* loss. Further studies may reveal insights on the mechanisms of CD44/SPP1 overexpression, functional significance in murine/ human renal tumorigenesis and utility as a prognostic marker or cell-surface target for ADC/ RPT-based therapeutic approaches in tRCC.

## Materials and Methods

### Cell lines

Patient-derived, human tRCC cell lines [*PRCC-TFE3* (UOK120 and UOK124) and *SFPQ-TFE3* (UOK145)] and HK-2 cells with stable, doxycycline-inducible expression of TFE3 proteins^30^ were a kind gift of Dr. W. Marston Linehan (NCI). <u>HEK293</u> cells with stable doxycycline-inducible expression of TFE3 proteins were generated using Flp Recombinase-mediated integration, using the Flp-In T-Rex Core Kit (K6500-01, Invitrogen). <u>UOK cells</u> were maintained in DMEM high glucose medium (11995065, Gibco) with l-glutamine, 10% heat-inactivated FBS (SH30071.03HI, Hyclone), 1X MEM (11130051, Gibco) and 1% penicillin/ streptomycin at 37°C in 5% CO2. <u>Inducible HEK cells</u> were maintained in DMEM high glucose medium, 10% FBS (Tet-approved; A4736401, Gibco), Blasticidin S (15 µg/ml) Hygromycin B (150 µg/ml) and 1% penicillin/ streptomycin. <u>Inducible HK-2 cells</u> were maintained in Advanced DMEM/F-12 (12634010, Thermo), 1.5% FBS (Tet-approved) 1X GlutaMAX (35050061, Thermo), Puromycin (0.8 µg/ml), Blasticidin S (2 µg/ml) and 1% penicillin/ streptomycin.

### Tissue samples

This study was approved by the Institutional Review Board of Johns Hopkins University and included four tissue microarray (TMA) cohorts, including the MiTF/TFE translocation renal cell carcinoma (TFE3 n=29; TFEB n=18), and the common subtypes: clear cell renal cell carcinoma (n=9) and papillary renal cell carcinoma (n=7)^31^. 2-5 redundant tumor samples and normal kidney tissue were included in each TMA.

### Immunohistochemistry and CD44 analysis

CD44 immunohistochemistry (IHC) on human tissues was performed using the Ventana Discovery ULTRA platform (Ventana/Roche, Oro Valley, AZ, USA) using hand-applied CD44 primary antibody (**E7K2Y** Rabbit mAb #37259, 1:100; Cell Signaling Technology, Danvers, MA, USA). TMA slides were scanned by Hamamatsu NanoZoomer S360 with NZAcquire and apparent magnification of 40x. Visual scores were determined by a trained urologic pathologist. The tumor intensity score, tumor proportion score, and tumor IHC score were visually graded using the criteria as previously described^32^.

### Animal Studies

Animal protocols were approved by the JHU Animal Care and Use Committee. The following strains were used: **1)** Mice hemizygous for the *<u>Ksp</u>*<u>-Cre recombinase</u> knockin gene (Strain Number: 012237) (The Jackson Laboratory), **2)** *PRCC-TFE3^LSL^* mice expressing the *PRCC-TFE3* fusion downstream of a *LoxP-Stop-LoxP (LSL)* cassette, were a kind gift of Dr. W. Marston Linehan (NCI)^23^, **3)** *SFPQ-TFE3^LSL^* mice expressing the *SFPQ-TFE3* fusion downstream of a *LoxP-Stop-LoxP (LSL)* cassette, were generated by Taconic Biosciences^21^. **4)** Tamoxifen-inducible, *Pax8* Cre-ER^T^^2^ mice were a kind gift of Dr. Athena Matakidou (Cancer Research, UK). Genomic DNA was isolated from tail snips and genotyping performed using primers, as previously described^21^.

### Histology and immunostaining

Mouse kidneys were fixed in 10% neutral buffered formalin (Sigma-Aldrich), embedded in paraffin, sectioned at 4 µm and used for H&E staining and immunohistochemistry.

### Immunoblotting

Cells and tissues were lysed in RIPA buffer supplemented with 10 μl Halt Protease and Phosphatase Inhibitor Cocktail (78440, Thermo Fisher Scientific). Lysates were centrifuged at 21,000 rpm for 10 minutes at 4°C and supernatants collected. Protein concentrations were quantified using the BCA Protein Assay Kit (23225, Pierce), and protein was resolved on 4-12% Bis-Tris SDS-PAGE gel (Thermo Fisher Scientific). Protein was transferred to nitrocellulose membranes (Amersham Bioscience), blocked for 1h at room temperature in 5% nonfat milk in 1X TBS-T and then incubated overnight with a primary antibody diluted in 5% milk in 1X TBS-T. The secondary antibodies used were anti-rabbit or anti-mouse immunoglobulin as appropriate (Cell Signaling) and diluted at 1:1000 in 5% nonfat milk in 1X TBS-T. Blots were developed using a chemiluminescent development solution (Super Signal West Femto, Pierce) and bands were imaged on a chemiluminescent imaging system (ChemiDoc Touch imaging System using the ImageLab Touch Software (version 2.3.0.07) (Bio-Rad). Digital images were quantified using Image J (version 1.52p) and all bands were normalized to their respective β-actin or GAPDH expression levels as loading controls.

### Plasmids, Lentiviral transfections and RNAi

Cells were transiently transfected using Lipofectamine RNAiMAX (13778075, Thermo; siRNA transfections) according to the transfection guidelines. CD44-targeting MISSION^®^ shRNA constructs were purchased as bacterial glycerol stocks (Sigma-Aldrich, SHCLNG -TRCN0000308110), while non-targeting control was purchased as plasmid (3rd gen lentiviral negative control vector containing scrambled shRNA, Addgene, Plasmid #1864) and introduced to competent cells.

### Antibodies and Reagents: Primary antibodies

**CD44** (37259 CST; 1:1000), **CD44 v6 (C44Mab-9)** (99618 CST; 1:1000), **Osteopontin/SPP1 (murine)** (88742 CST; 1:1000), **Osteopontin/SPP1 monoclonal antibody (7C5H12) (human)** (MA5-17180 Thermo Fisher; 1:1000), **GPNMB** (90205 CST; 1:1000), **GPNMB** (38313 CST; 1:1000), **TFE3** (ABE1400 Sigma; 1:4000), **GAPDH** (5174 CST; 1:1000), **Phospho-S6 Ribosomal Protein (Ser235/236)** (4858 CST; 1:2000), **Phospho-4E BP1 (Ser65)** (9451 CST; 1:2000), **Phospho-4E BP1 (Thr37/46)** (2855 CST; 1:1000), **Phospho-p70 S6 Kinase (Thr389)** (9234 CST; 1:1000).

### RNA sequencing and data analysis

RNA sequencing of quadruple UOK cell line replicates was performed at Plasmidsaurus and carried out as previously described^33^. Raw RNAseq counts were Fragments Per Kilobase of transcript per Million mapped reads (FPKM) or Transcripts Per Kilobase Million (TPM)-normalized for data visualization in R (v4.3.2). Raw counts were also imputed in DESeq2 in R to determine differentially expressed genes. Log2 fold-changes, p-values, and adjusted p-values (false discovery rate method, FDR) were obtained for all genes and comparisons. Processed RNAseq data from the following cohorts was accessed in GEO: 1) ***Sun et al*** (GSE167573), 2) ***Wang et al*** (GSE150474), 3) ***Bakouny et al*** (GSE188885), 4) ***Prakasam et al*** (GSE252047) and 5) ***Baba et al*** (GSE130072). Normalized gene expression values for *Cd44* and *Spp1* were extracted and quantified as FPKM or TPM. Data visualization was performed using R (via the *ggplot2* package), with expression distributions represented as boxplots overlaid with jittered individual data points. Statistical differences in FPKM values between the respective transgenic fusion models and their wild-type/control counterparts were evaluated using two-tailed unpaired Student’s t-tests. Statistical significance was defined as *p* < 0.05. Pan-Cancer-normalized RNAseq data from TCGA was downloaded from TCGA Pan-Cancer publication portal (https://gdc.cancer.gov/about-data/publications/panimmune). KIRP, KIRC and KICH RNAseq data were previously normalized with the methods Fragments Per Kilobase of transcript per Million mapped reads (FPKM) and FPKM Upper Quartile (FPKM-UQ) by the TCGA research team. Gene expression data was compared using Wilcoxon rank sum tests adjusted with multiple comparisons, where applicable. GSEA analyses were conducted with raw RNAseq counts using custom pathways and standard enrichment molecular signatures database. All TCGA graphs presented in the manuscript were generated by us using TCGA data. All analyses were performed in R v4.3.1.

### Single-nuclei RNA-Seq analysis

We subjected kidney tissue from SFPQ–TFE3 fusion transgenic mice to single-nucleus RNA-seq using Chromium GEM-X Single Cell 3′ v4 (10x Genomics). Samples were obtained from mice fifteen days or six months after tamoxifen treatment, as well as the respective time-matched untreated controls. Sequenced reads were mapped to a GRCm39-derived splici reference using simpleaf^34^. The resulting count matrices were then imported into CellBender for removal of ambient RNA^35^. Preprocessed data were imported into Scanpy for quality control and downstream analysis^36^. We retained nuclei with more than 750 detected genes and more than 750 UMIs, with mitochondrial counts comprising no more than 2.5% of total counts. Counts were depth-normalized and log-transformed, and nuclei were clustered using igraph’s implementation of Leiden algorithm^37,38^ on a nearest-neighbor graph constructed from principal components harmonized across samples using Harmony^39^. Principal component analysis was based on a subset of 4,000 highly variable genes identified using Scanpy’s implementation of the Seurat v3 method^40^. Cell-type annotation was performed in the harmonized space by inspecting Leiden clusters for markers identified using Presto^41^, ranked by detection-frequency difference, auROC, and difference in mean log-normalized expression. The resulting cell-type marker lists were cross-checked and refined with reference to previously published studies^42–44^. To visualize sample-specific differences, we plotted gene expression on the non-harmonized UMAP embedding (DOI: 10.21105/joss.00861), alongside sample IDs and cell types.

## Acknowledgements

This research was supported in part by the CDMRP TSCRP grant W81XWH-22-1-0264 (TLL), CDMRP KCRP grant W81XWH-20-1-0843 (TLL), CDMRP KCRP grant W81XWH2210377 (KA), KCA-2024-TRANS-001 (Kidney Cancer Association Translocation Award (KA)) and the NCI Cancer Center Support Grant 5P30CA006973-52. This work was supported at Johns Hopkins in part by Dahan Translocation Carcinoma Fund and Joey’s Wings Foundation. This research was supported by the Intramural Research Program of the NIH, National Cancer Institute, Center for Cancer Research (WML). This project has been funded in whole or in part with Federal funds from the National Cancer Institute, National Institutes of Health, under Contract No. HHSN261201500003I. (LSS). The content of this publication does not necessarily reflect the views or policies of the Department of Health and Human Services, nor does mention of trade names, commercial products, or organizations imply endorsement by the U.S. Government.

## Contributions

K.A. conceived the study. K.A. drafted the manuscript. K.A., J.W., T.V., K.F., A.L., R.S., A.T., S.R., H.L., L.O., A.A., V.P., R.A., J.P., E.I., R.E., C.T., H.G., A.G., P.P., J.D., Y.G., N.S., J.C., L.S.S., W.M.L., P.A., T.L.L. completed data collection and analysis. All authors critically reviewed the manuscript and agreed to submit it for publication.

